# Metabolic flux signatures of the ER unfolded protein response *in vivo* reveal decreased hepatic *de novo* lipogenesis and mobilization of lipids from adipose tissue to liver

**DOI:** 10.1101/2020.10.29.360073

**Authors:** Catherine P. Schneider, Lucy Peng, Samuel Yuen, Michael Chang, Rozalina Karapetyan, Edna Nyangau, Hussein Mohammed, Hector Palacios, Naveed Ziari, Larry K. Joe, Ashley E. Frakes, Andrew Dillin, Marc K. Hellerstein

## Abstract

The unfolded protein response in the endoplasmic reticulum (UPR^ER^) is involved in a number of metabolic diseases, including non-alcoholic fatty liver disease. Here, we characterize the UPR^ER^ induced metabolic changes in mouse liver through *in vivo* metabolic labeling and mass spectrometric analysis of proteome and lipid fluxes. We induced ER stress *in vivo* via tunicamycin treatment and measured rates of proteome-wide protein synthesis, *de novo* lipogenesis and cholesterol synthesis serially over a three-day period, thereby generating a metabolic “signature” of the UPR^ER^ over time. Synthesis of most proteins was suppressed under ER stress conditions, including proteins involved in lipogenesis, consistent with reduced *de novo* lipogenesis at 48 and 72 hours. Electron microscopy revealed striking morphological changes to ER and H&E staining showed lipid droplet enriched livers under ER stress. Pre-labeling of adipose tissue prior to ER stress induction revealed mobilization of lipids from adipose to the liver. Interestingly, the source of these lipids was uptake of free fatty acids, not whole triglycerides or phospholipids from lipoproteins, as demonstrated by replacement of the triglyceride-glycerol moiety in liver concurrently with increased incorporation of labeled palmitate from adipose. We also induced ER stress by a high-fat diet and observed similar metabolic flux signatures, suggesting that this mechanism may play a role in the progression of fatty liver disease. This flux-based approach provides a powerful tool to identify novel regulators of ER stress and potential targets for pharmacological intervention.

## Main

Proteostasis, or protein homeostasis, is important in maintaining a healthy cellular environment under stressful conditions^1^. Dietary changes such as increased intake of fatty acids or the accumulation of misfolded proteins can perturb proteostasis and thereby initiate the endoplasmic reticulum unfolded protein response (UPR^ER^)^2–4^. The role of the UPR^ER^ in metabolic diseases such as non-alcoholic fatty liver disease has been hypothesized to be due to dysregulation of lipid homeostasis but remains poorly understood^5–8^. Here, our aim was to characterize metabolic flux changes of UPR^ER^, including rates and metabolic sources of lipid synthesis as well as protein synthesis rates across the proteome.

The UPR^ER^ consists of three arms, led by ER membrane anchored IRE1L, PERK, and ATF6. In times of ER stress, BiP, a key chaperone, moves away from the ER membrane to combat the accumulation of misfolded protein and activates these three arms^9^. Hallmarks of the UPR^ER^ include suppressed global protein translation, with the exception of key ER stress responders such as chaperones^10^. Although these responses have been characterized at the mRNA and protein concentration level, rates of protein synthesis (translation rates) in response to induction of the ER stress response *in vivo* have yet to be characterized. Here we used heavy water labeling and proteome-wide tandem mass spectrometric analysis to measure newly synthesized proteins after tunicamycin-induced ER stress in mice. Because synthesis of most protein is suppressed under ER stress conditions, identifying any exceptions is a potentially powerful tool to identify novel features or regulators of ER stress. Translation rate also may shift over time, and which proteins are synthesized at different points after ER stress induction has not yet been established.

Another hallmark of the UPR^ER^ is thought to be an increase of lipogenesis by a stressed cell to explain the expansion of ER lipids. This has been studied mostly in cell culture models^11^. Controversy about this canonical pathway exists, however, as it has been shown *in vivo* that lipogenic gene expression is reduced in livers of mice under ER stress^12,13^. Increase in lipogenesis under ER stress conditions may be important in providing added surface for resolution of ER protein synthetic stress through re-folding and clearance of misfolded proteins in the ER lumen^14^, but other metabolic sources in a whole organism could be responsible for increases in ER lipids, including import from extracellular sources and other tissues.

We were curious how lipogenesis rates change over time after ER stress induction *in vivo* and how this might be integrated with changes in protein translation rates. We used heavy water labeling to measure rates of *de novo* lipogenesis, cholesterol synthesis, and protein synthesis rates across the proteome in the liver over a three-day labeling period after tunicamycin-induced ER stress. We performed RNA-seq on the liver tissue to compare mRNA levels to rates of protein synthesis and lipogenesis. We report here a decline in *de novo* lipogenesis by all these metrics: reduced *de novo* lipogenesis and cholesterol synthesis flux rates, reduced synthesis rates of lipogenic proteins, and reduced expression of lipid-synthesis related genes. Even so, electron microscopy and hematoxylin and eosin staining visualized lipid accumulation and changes in ER membrane morphology over this time course. To explain the metabolic source of hepatic lipid accumulation, we then labeled adipose triglycerides by long-term heavy water administration and allowed body water label to die-away prior to ER stress induction. We demonstrate that the lipids incorporated into ER membranes or that accumulate in lipid droplets during ER stress in the liver *in vivo* are mobilized from other tissues.

## Results

### RNA-seq of mouse liver under induced ER stress reveals decreased expression of genes involved in lipid and cholesterol synthesis

To investigate what happens over the time following initiation of the UPR^ER^, we used tunicamycin to induce ER stress in mice and took liver samples over the subsequent three days (figure 1a). RNA-seq of the liver tissue revealed many significant changes, including decreased expression of genes involved in lipid synthesis, cholesterol metabolism, and glutathione synthesis. Upregulated ontologies included genes involved in response to ER stress, ERAD, and ribosome biogenesis (figure 1b-g). Expression of genes involved in the UPR^ER^ shift over time post ER-stress induction (figure 1b-f).

**Figure 1:**
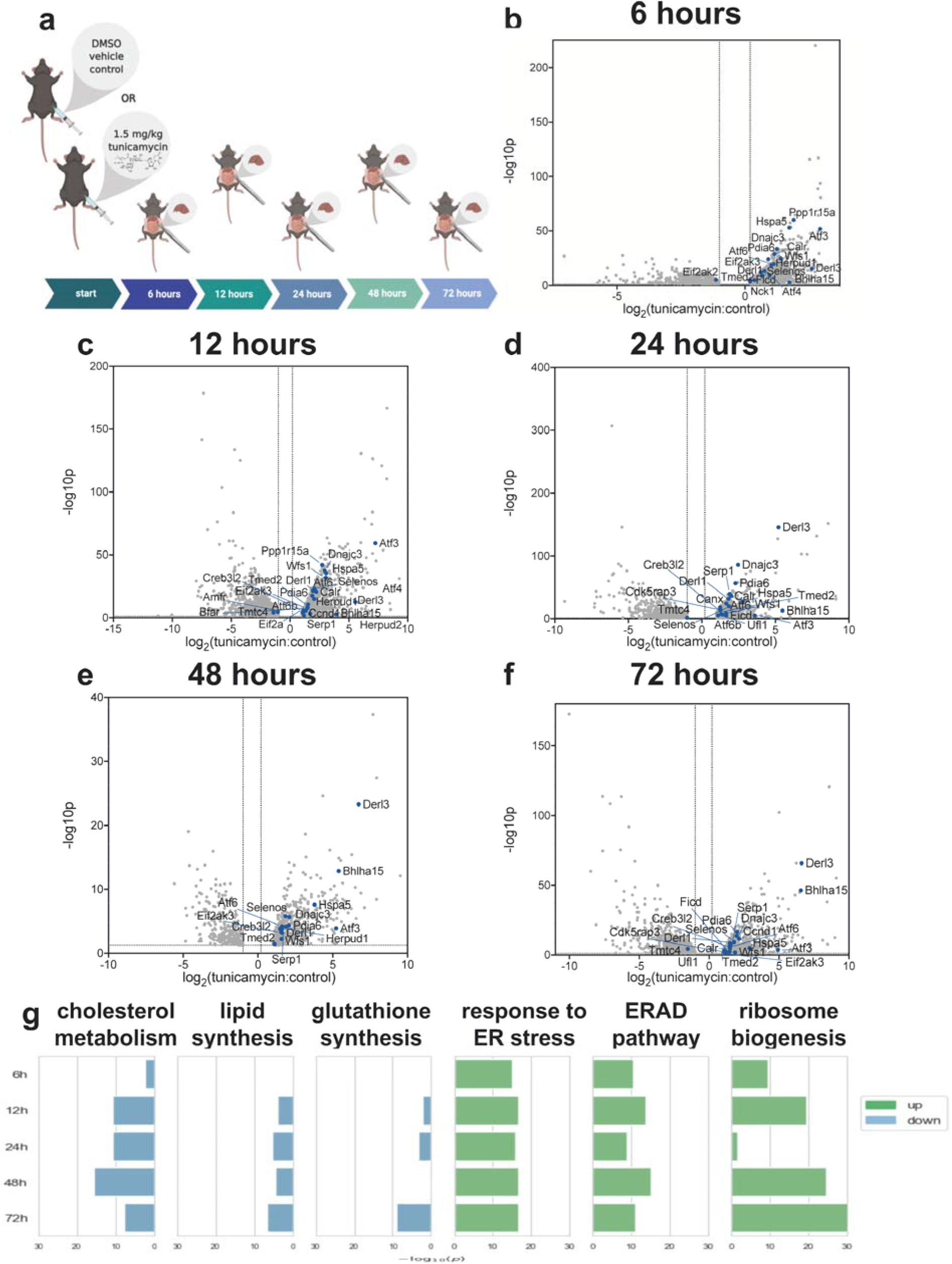
(a) Experimental overview. Mice (n=5 per group) were treated with either DMSO or 1.5 mg/kg tunicamycin. Tissues were taken 6-72 hours post-treatment. (b-f) Volcano plot of all genes for which RNA-seq measured expression. Points expressed as log2 fold-change tunicamycin treated/control on x-axis and − log10(p-value), obtained from 2-tail t-test, on y-axis. (g) GO (gene ontology) analysis for genes for which tunicamycin treatment significantly changed gene expression. GO-terms indicate groups for which a significant number of genes where changed in relation to the remainder of the data set for each time point. GO with significant decreased gene expression: cholesterol metabolic process (GO: 0008203), lipid biosynthetic process (GO: 0008610), glutathione biosynthetic process (GO: 0006749). GO with significant increased gene expression: response to ER stress (GO: 0034976), ERAD pathway (GO: 0036503), ribosome biogenesis (GO: 0042254).

### Dynamic proteomics measurements reveal decreased global protein synthesis rates, including those involved in lipid synthesis but not key UPR^ER^ proteins

We asked whether protein translation rates would match the trends identified via RNA-seq using the dynamic proteomic approach^15^ to measure translation rates of proteins after UPR^ER^ induction (figure S1a). Deuterated water was administered concurrently with tunicamycin to label nascent proteins that were translated over the three-day treatment period. During the first 12 hours post-tunicamycin, synthesis rates of most proteins measured were suppressed, with the exception of proteins involved in ER stress, including BiP, protein disulfide isomerases, and other chaperones (figures 2a-f). Proteins for which translation rates were significantly increased or decreased with induction of ER stress as compared to controls were organized into KEGG-pathways for analysis (figure 2g). Synthesis of proteins involved in fatty acid metabolism were decreased starting at 6 hours post tunicamycin treatment (figure 2g), as were most proteins characterized as being involved in “metabolic pathways”. By 12 hours post treatment, some protein translation rates began to recover, while others remained reduced through 48 hours.

**Figure 2:**
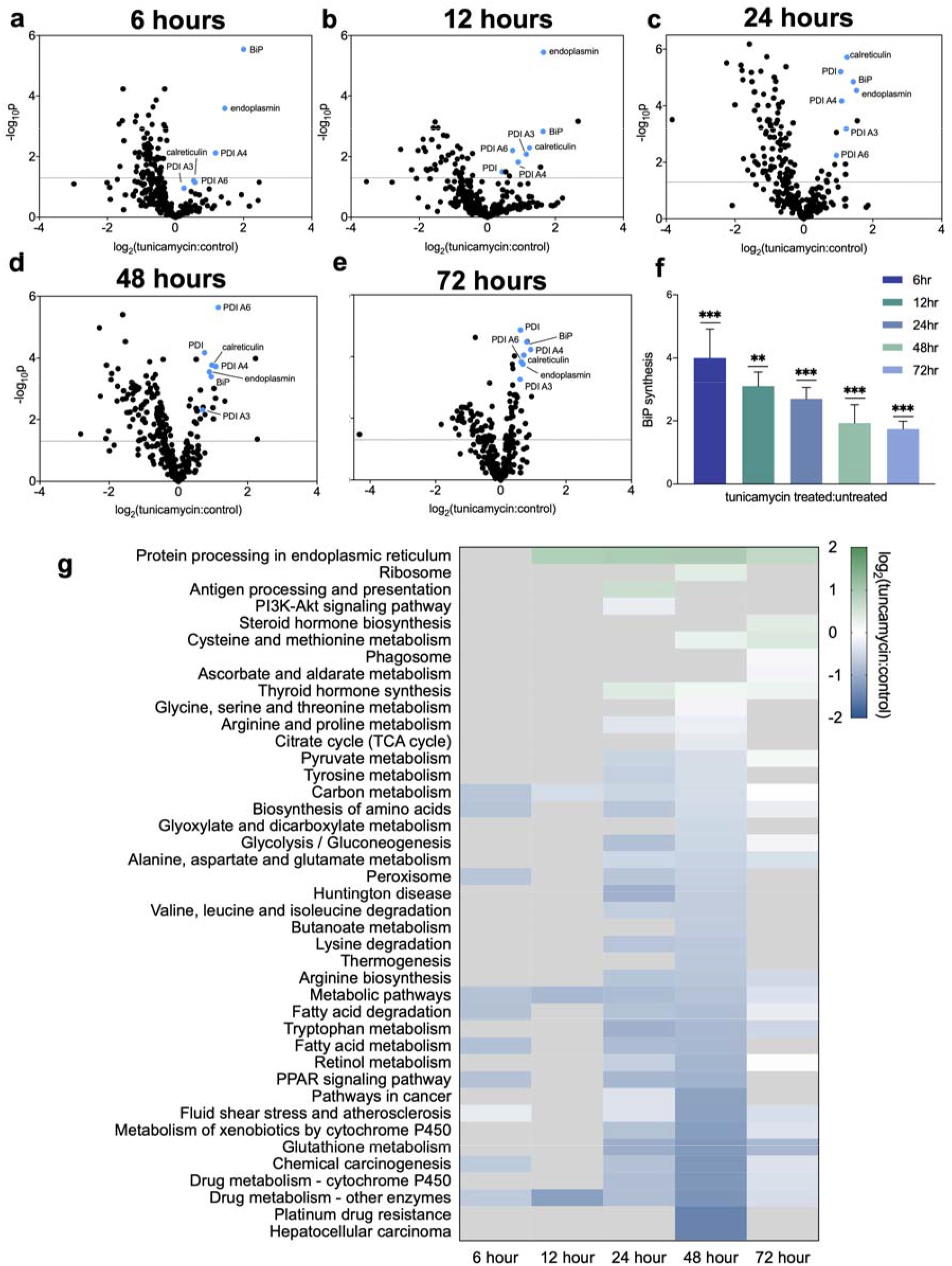
(a-f) Volcano plot of all proteins for which fractional synthesis rates were measured 12-72 hrs post tunicamycin treatment. Points expressed as log2 fold-change tunicamycin treated/control on x-axis and − log10(p-value), obtained from 2-tail t-test, on y-axis. (f) ratio of protein translation rates of BiP in tunicamycin treated/control. ns= no significance, * = <0.05, ** = <0.01, *** = <0.001. (g) KEGG-pathway analysis for fractional synthesis rates of significant (p=<0.05 per 2-tail t-test) proteins from tunicamycin treated/control. n=at least 5 proteins per pathway.

Most protein synthesis rates returned to baseline values at the final time point measured, 72 hours (figure 2g). KEGG-pathway analysis revealed significantly increased synthesis rates of proteins involved in protein processing in the ER from the 12-hour time point continuing to the 72-hour time point, which can be attributed to the many chaperones and other UPR^ER^ responders. In particular, BiP showed a marked increase in synthesis rate at earlier time points, and though trending down over time, remained elevated at 72 hours (figure 2c). KEGG pathway analysis showed markedly decreased synthesis rates of proteins involved in lipid metabolism at 24 and 48 hours (figure 2g). Decreased synthesis of proteins involved in glutathione synthesis matched the decline observed in the RNA-seq data (figure 1b, figure 2g). Proteins involved in ribosomal biogenesis were also increased at the 48-hour timepoint, matching the RNA-seq data (figure 1b, figure 2g).

### Changes in ER morphology by electron microscopy

We asked how the decline of gene expression and protein synthesis rates of lipid synthesis proteins correlated with ER membrane expansion reported by other groups under ER stress conditions^14^. Electron microscopy was carried out to visualize liver taken at each time point over the three-day treatment period. We observed distinct morphological changes to ER structure beginning at 12 hours post-tunicamycin treatment and continuing through the 72-hour endpoint. ER in the control samples presented as stacked and ribosome studded, as expected, whereas the ER observed in the tunicamycin-treated animals appeared like bubbles, or cobblestone, and appeared to be barren of the usual ribosomes (figure 3a).

**Figure 3:**
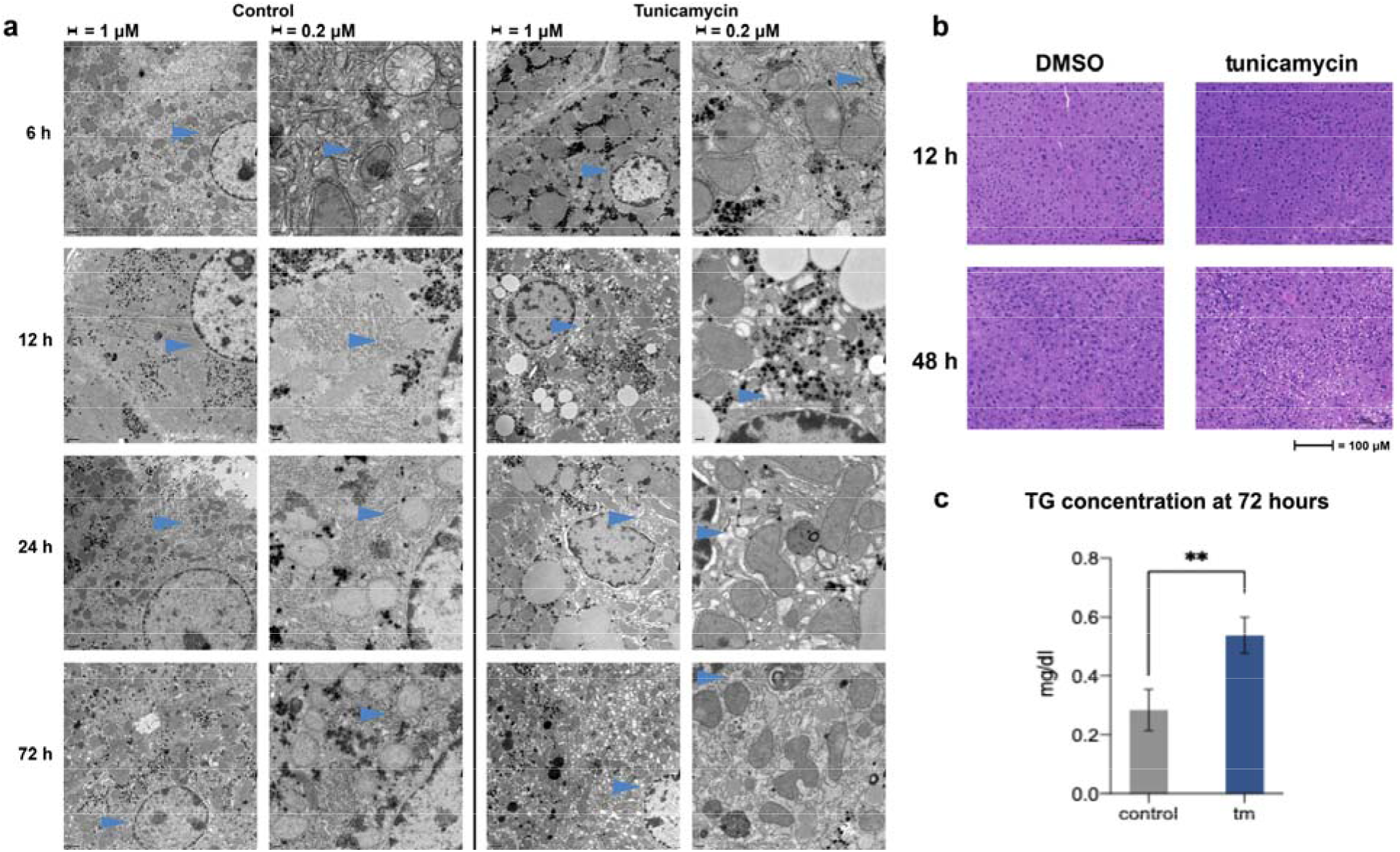
(a) Electron microscopy (TEM) images of liver sections 12-72 hours post DMSO (control) or 1.5 mg/kg tunicamycin treatment. Arrows point out ER in both treated and control to highlight changes in morphology. (b) H&E staining of liver sections 12 and 48 hrs post DMSO or 1.5 mg/kg tunicamycin treatment. (c) Concentration of triglycerides in mouse liver post DMSO (control) or 1.5 mg/kg tunicamycin treatment. (Tm=tunicamycin-treated).

### ER stress induced lipid accumulation by histology

Hematoxylin and eosin (H&E) staining of liver taken from mice treated with tunicamycin or DMSO revealed lipid accumulation in the liver starting at 48-hours post ER stress induction. Earlier time points revealed no lipid differences as compared to controls (figure 3b).

### Lipid and cholesterol synthesis rates decreased at later time points post-ER stress induction

Due to the striking morphological changes, we asked if lipids in the newly expanded ER membrane and droplets were coming from *de novo* synthesis. This was measured from the heavy water labeling of fatty acids in phospholipids and triglycerides and in free and esterified cholesterol (figure S1b)^16^. The contribution from *de novo* lipogenesis to palmitate in hepatic triglycerides and phospholipids was significantly decreased in both fractions beginning at 48-hours post tunicamycin treatment and continued through the last time point at 72-hours (figure 4a-b). *De novo* synthesis of both free and esterified cholesterol in the liver was also significantly decreased at 48 and 72-hours post ER stress initiation (figure 4c), consistent with the significant decline in expression of cholesterol synthesis genes in liver.

**Figure 4:**
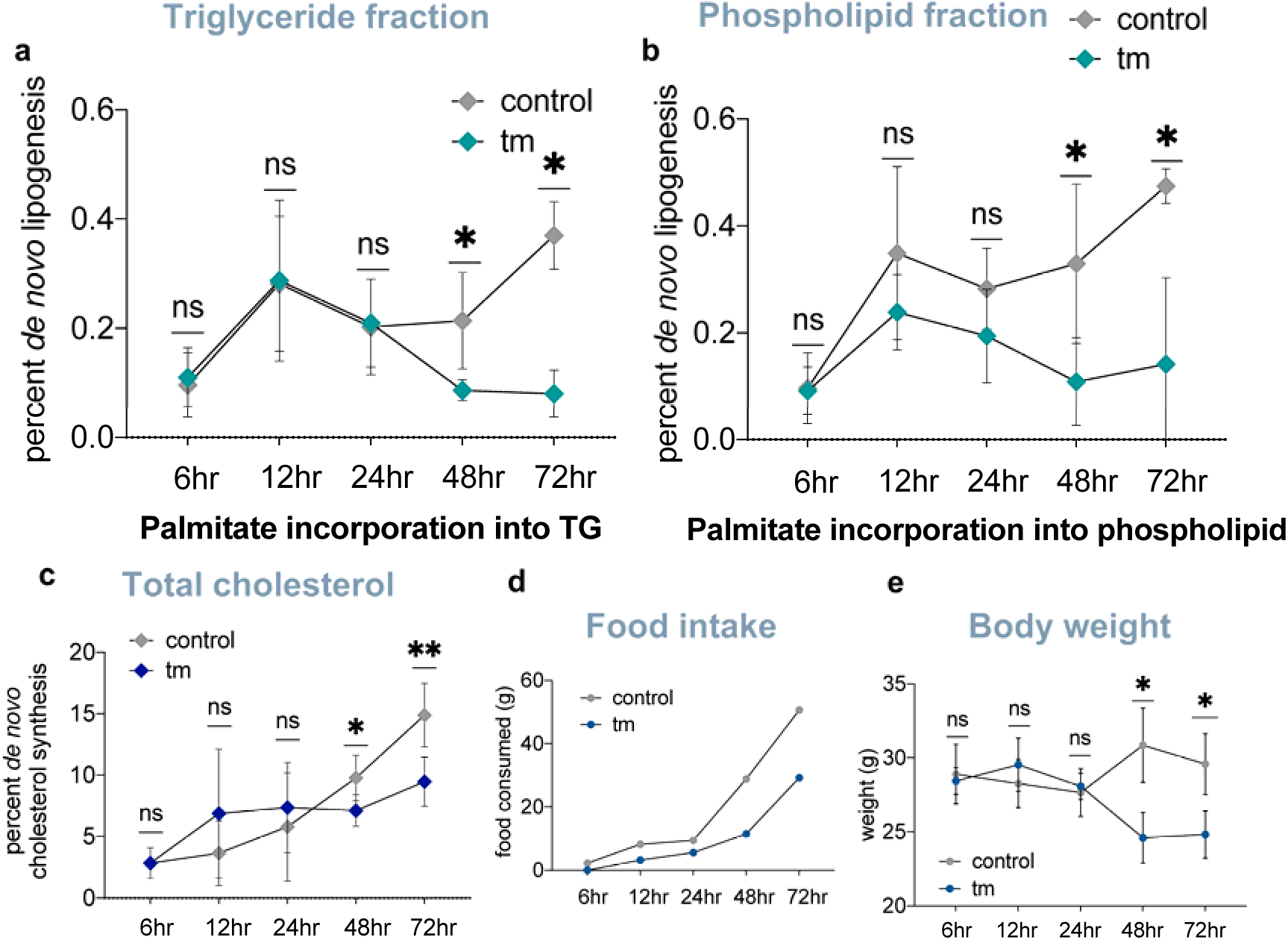
(a) *De novo* lipogenesis rates of palmitate incorporated into triglycerides in control and tunicamycin treated mice 12-72 hours post treatment (n=5 per group). (b) *De novo* lipogenesis rates of palmitate incorporated into phospholipids in control and tunicamycin treated mice 12-72 hours post treatment (n=5 per group). (c) *De novo* cholesterol synthesis (free and esterified) rates in control and tunicamycin treated mice 12-72 hours post treatment (n=5 per group). (d) Food intake in mice control and tunicamycin 12-72 hours post treatment (n=5 per group). (e) Average body weight of control and tunicamycin treated mice 12-72 hours post treatment (n=5 per group). ns= no significance, * = <0.05, ** = <0.01, *** = <0.001.

### Mobilization of lipids from adipose tissue to liver under ER stress conditions

To answer the question of where lipids in the liver were coming from if not from *de novo* synthesis, we devised a protocol to pre-label extrahepatic triglycerides (i.e., primarilly adipose tissue lipid stores) prior to inducing ER stress. Mice were given deuterated water for 7 weeks to incorporate newly synthesized fatty acids into adipose tissue. We then discontinued heavy water intake for 2 weeks to allow deuterium enrichment to die away in body water and in liver triglycerides, which have much shorter half-lives than triglycerides in adipose tissue^17^. (figure 5a-b). After tunicamycin-induced ER stress, deuterium enrichment of palmitate in hepatic triglycerides increased whereas deuterium enrichment of palmitate in adipose tissue triglycerides decreased compared to controls (figure 5c), consistent with lipid mobilization from adipose tissue to the liver. Additionally, we observed a reduction in enrichment of glycerol in phospholipids in the liver, indicating that intact phospholipids were rapidly turned over to the free glycerol level under ER stress conditions (figure 5d). A decline in triglyceride-glycerol enrichment in the liver concurrently with increased palmitate enrichment indicates that pre-existing hepatic triglycerides were hydrolyzed free fatty acids and free glycerol, not to the level of mono- or di-glycerides, and that the influx of palmitate is in the form of free fatty acids as opposed to transport of whole triglycerides or phospholipids from lipoproteins (figure 5c-d).

**Figure 5:**
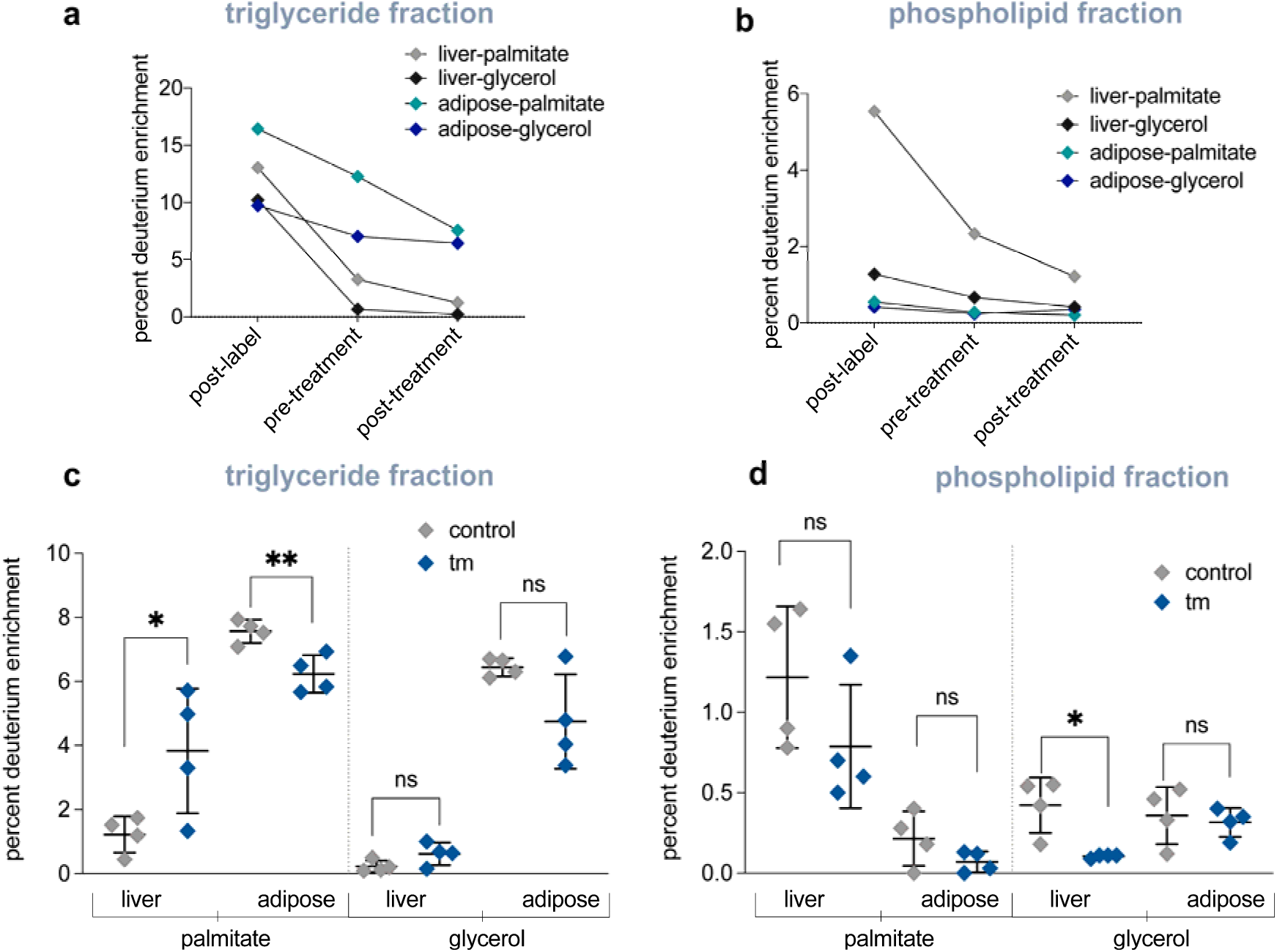
(a) Pre-labeling of triglycerides in control (DMSO) mice: percent deuterium enrichment of palmitate incorporated into triglycerides after 7 weeks of deuterium labeling (post-label), after a 2-week label free period (pre-treatment), and post 72-hour treatment period (n=3 per group). (b) Pre-labeling of phospholipids in control (DMSO) mice: percent deuterium enrichment of palmitate incorporated into phospholipids after 7 weeks of deuterium labeling (post-label), after a 2-week label free period (pre-treatment), and post 72-hour treatment period (n=3 per group). (c) Percent deuterium incorporated into palmitate and glycerol of triglycerides found in the liver or adipose after control (DMSO) or 1.5 mg/kg tunicamycin treatment after 72 hours (n=4 per group). (d) Percent deuterium incorporated into palmitate and glycerol of phospholipids found in the liver or adipose after control (DMSO) or 1.5 mg/kg tunicamycin treatment after 72 hours (n=4 per group). ns= no significance, * = <0.05, ** = <0.01, *** = <0.001.

### Diet induced changes to protein and lipid metabolic flux signatures

To understand if this metabolic signature of was conserved under other models of UPR^ER^, such as lipotoxicity, we used prolonged high-fat diet to induce ER stress *in vivo* (figure 6a), and also combined high-fat diet with tunicamycin treatment to determine their additive effects on the ER stress response. High-fat diet alone led to an overall increase in translation rates proteome wide, with many canonical UPR^ER^ proteins being significantly upregulated in their translation rates (figure 6c-d). We also saw a decrease in *de novo* lipogenesis rates in both triglyceride and phospholipid fractions through high-fat diet induced ER stress (figure 6b). When high-fat diet was coupled with tunicamycin stress induction, protein synthesis rates were mostly suppressed with the exception of canonical UPR^ER^ proteins (figure 6e-f), and an exaggerated decrease in *de novo* lipogenesis rates in both triglyceride and phospholipid fractions was seen (figure 6g).

**Figure 6:**
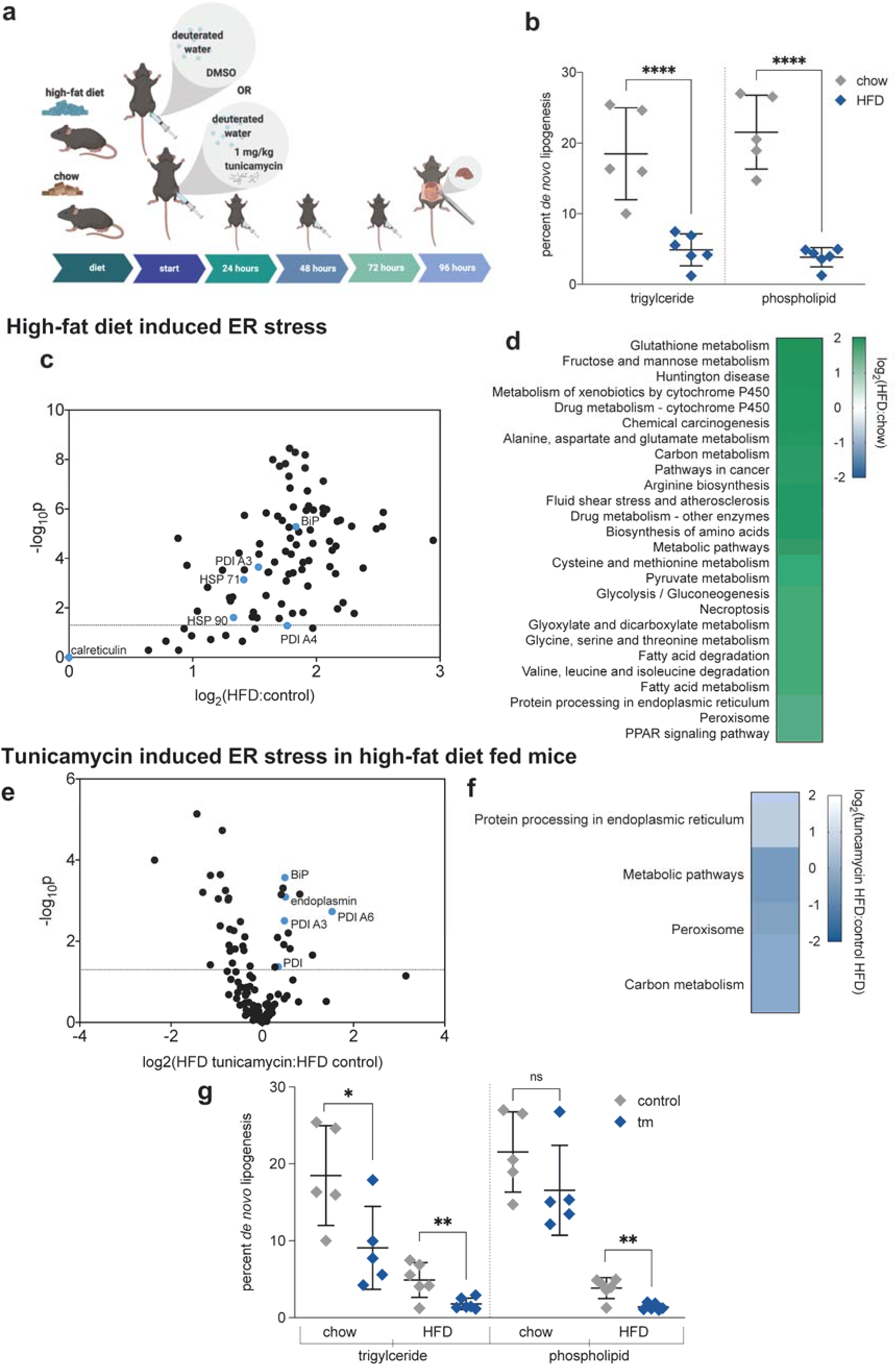
High-fat diet induced ER stress and effect of high-fat diet on ability to handle additive tunicamycin induced ER stress. All data from mouse livers. (a) Experimental overview: mice were given high-fat or chow diet for 6 weeks. Mice were challenged with a dose of 1 mg/kg tunicamycin daily until samples were taken at 96-hours post initial treatment. (b) Percent *de novo* lipogenesis of palmitate in high-fat diet compared to chow fed mice in both triglycerides and phospholipids. (c) Volcano plot of all proteins for which fractional synthesis rates were measured 96 hours after deuterium labeling in either chow or high-fat diet fed mice. Points expressed as log2 fold-change high-fat diet/chow on x-axis and − log10(p-value), obtained from 2-tail t-test, on y-axis. (d) KEGG-pathway analysis for fractional synthesis rates of significant (p=<0.05 per 2-tail t-test) proteins from high-fat diet/chow. n=at least 5 proteins per pathway. (e) Volcano plot of all proteins for which fractional synthesis rates were measured 96 hours after tunicamycin treatment in high-fat diet fed mice. Points expressed as log2 fold-change tunicamycin/control on x-axis and − log10(p-value), obtained from 2-tail t-test, on y-axis. (f) KEGG-pathway analysis for fractional synthesis rates of significant (p=<0.05 per 2-tail t-test) proteins from tunicamycin/control. n=at least 5 proteins per pathway. (g) Percent *de novo* lipogenesis of palmitate in tunicamycin compared to control treated high-fat diet fed mice in both triglycerides and phospholipids. n=6 mice per group. ns= no significance, * = <0.05, ** = <0.01, *** = <0.001.

**Figure 7:**
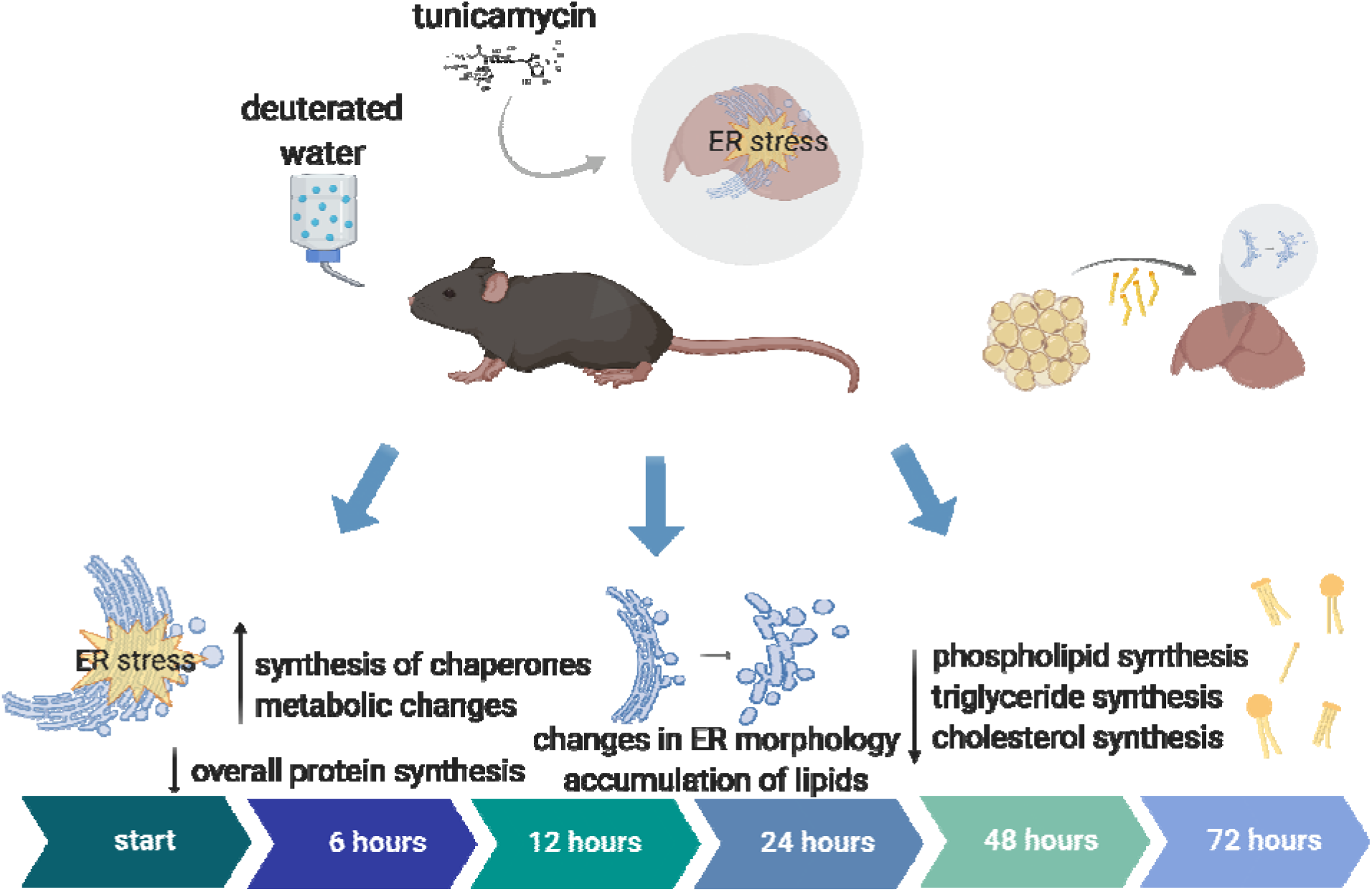
Summary figure shows metabolic flux signatures post tunicamycin induced ER stress in mouse liver. Overall protein synthesis declines with the exception of chaperones and UPR^ER^ related proteins. Proteins involved in lipogenesis are significantly decreased in their synthesis rates. At 12 hours post tunicamycin, ER stress induced changes in ER morphology in hepatocytes are evident, followed by lipid accumulation at 48 hours. At 48 hours post ER stress induction, palmitate incorporated into both phospholipids and triglycerides decline, as well as cholesterol synthesis. Lipids appear to be mobilized from the adipose tissue and deposited in the liver.

## Discussion

Metabolic responses to the initiation of the UPR^ER^ are not well understood^18^. We used metabolic labeling with stable isotopes to concurrently measure rates of fatty acid, cholesterol, and protein synthesis after inducing ER stress, with the goal of characterizing metabolic flux signatures over time and their relation to ultrastructural changes and gene expression patterns. Overall, proteome-wide protein synthesis rates declined with tunicamycin-induced ER stress, with the exception of chaperones and other key ER proteins recognized to be induced during the UPR^ER19^. Protein synthesis rates generally matched the signatures measured through RNA-seq, which were similar to canonical UPR^ER^ signatures reported previously^20,21^. These data support the validity of using dynamic proteomics, as we have described previously^15,22^, as a method to study tunicamycin-induced ER stress. This signature appears to be unique to the liver, as kidney tissues from the same tunicamycin treated mice failed to present a similar protein synthesis signature (figure S2). In other physiologic conditions, in contrast, clear dissociation between mRNA and protein synthesis rates has been observed^23–25^. Measurement of protein fluxes provides a potentially powerful tool for identifying UPR^ER^ regulators and signatures.

In particular, we found that synthesis rates of hepatic proteins involved in lipid metabolism and cholesterol synthesis were decreased in response to tunicamycin. In combination with similar reductions in gene expression for lipogenic proteins, this led us to investigate in depth *de novo* synthesis rates of lipids (palmitate in triglycerides and phospholipids) and cholesterol (free and esterified). These kinetic signatures somewhat surprisingly revealed significant decreases for both lipid and cholesterol synthesis rates over 72 hours under ER stress conditions.

These reductions in *de novo* lipid synthesis rates, gene expression, and protein synthesis rates for lipogenic enzymes were particularly striking in view of the well established changes in ER membrane structure and lipid stores during the UPR^ER11^. Our ultrastructural observations confirmed that after 12 hours, ER membranes appeared strikingly different by electron microscopy. The ER appeared almost bubble-like, consistent with expansion believed to create space for chaperone refolding of accumulated misfolded proteins as part of the adaptation to restore protein homeostasis^14,26–28^. H&E staining also showed that lipids accumulated in the liver at 48 hours, the same time point at which *de novo* lipid synthesis rates were significantly decreased. These results in living mice differ from some studies in isolated cells^11,29,30^, which have reported an increase in expression of genes involved in *de novo* synthesis during the UPR^ER^. We then demonstrated directly that new lipids in the liver *in vivo* are mobilized from other tissues such as those in adipose stores. Interestingly, the biochemical form of this lipid import appears to be as free fatty acids, not transport of intact triglyceride or phospholipids in plasma lipoproteins, based on the replacement of pre-labeled glycerol moiety of hepatic triglycerides concurrent with increased palmitate import form adipose tissue. Tunicamycin induced anorexia, as previously characterized by other groups^12^, was observed in our studies. Mice treated with tunicamycin ate and weighed less than control mice, and presented with less adipose tissue upon dissection, supporting the observed mobilization of lipids from adipose to the liver.

To evaluate these findings in a model more physiologically relevant to metabolic diseases, we also used a high-fat diet to induce ER stress through lipotoxicity and applied the same kinetic metrics to develop a metabolic flux signature^31^. Under a high-fat diet alone, most protein synthesis rates increased rather than decreased, including significant increases in canonical UPR^ER^ with similar increases to our findings with tunicamycin-induced ER stress. Lipid bilayer stress is thought to act though IRE1□ sensors, initiating downstream effects of this arm of the UPR^ER32^. Because inhibition of translation occurs downstream of the PERK arm of the UPR^ER^, it is reasonable to conclude that we did not see the same signature of proteome-wide suppression of protein synthesis in this lipotoxicity ER stress model as compared to the proteotoxicity model induced by tunicamycin. Other groups have also reported that a short term high-fat diet increases rather than decreases rates of protein translation^33,34^. We also saw decreased *de novo* lipogenesis rates under high-fat diet induced ER stress, which were exaggerated by the addition of tunicamycin induced ER stress. Tunicamycin also counteracted the widespread increases in protein synthesis rates induced by high-fat diet alone. We believe that this model demonstrates that livers under stress induced by high amounts of exogenous lipids exhibit a mixed handling of ER stress, which may not be as protective as with proteostatic stress alone and may be less able to restore normal homeostasis. This may further exacerbate a disease phenotype such as non-alcoholic fatty liver disease.

In summary, these data extend the metabolic alterations invoked during the UPR^ER^ and their importance in metabolic homeostasis. The source of accumulated lipid droplets and ER lipids during ER stress in the liver *in vivo* is from lipids taken up from outside the liver not synthesized *de novo* locally. Under ER stress conditions, key metabolic pathways including lipid and cholesterol synthesis are reduced while other pathways are perturbed in complex and not entirely predictable ways, including mobilization of lipids from adipose tissue to the liver. Prolonged disruption of these pathways may lead to progression of diseases involving altered lipid and protein homeostasis such as non-alcoholic fatty liver disease. This finding is useful as a differential pathogenic signature of ER stress in contrast to insulin-induced lipid accumulation in the liver, for example, where *de novo* lipogenesis is highly elevated^35^. These findings support that the UPR^ER^ may have implications for metabolic diseases characterized by accumulation of lipids.

## 4 Methods

### 4.1 Animals

C57BL/6J mice acquired from The Jackson Laboratory were used for this study. Mice were 12 weeks of age. All mice were housed according to the Animal Care and Use Committee (ACUC) standards in the animal facility at UC Berkeley. Mice were fed a standard chow diet and water ad libitum. Mice on a high-fat diet were fed a 60% high-fat diet (Research Diets, D12492) for 6 weeks.

### 4.2 Deuterated water labeling and tunicamycin treatment in mice

Mice were labeled with deuterated water (heavy water, ^2^H_2_O) beginning at time point 0 (t^0^) through the end of the experiment. Proteins synthesized after t^0^ will incorporate deuterium-labeled amino acids, thus enabling the measurement of proteins synthesized during the period of exposure to heavy water. Deuterium is rapidly incorporated throughout the body of an organism after treatment, bringing the deuterium enrichment in body water up to 5%. Deuterium enrichment is maintained through the intake of 8% ^2^H_2_O given as drinking water, thus making it an optimal labeling approach for *in vivo* experimental study. Mice are injected intraperitoneally (IP) with 100% ^2^H_2_O containing either tunicamycin dissolved in DMSO, or DMSO control. Mice were treated with 1.5 mg/kg tunicamycin dissolved in DMSO at t^0^, or DMSO control, and tissues were harvested 6, 12, 24, 48, and 72 hours after the initial injection (n=5 mice per group).

### 4.3 Deuterated water labeling and tunicamycin treatment in mice: pre-label of adipose tissue triglycerides

Mice were labeled with deuterated water (heavy water, ^2^H_2_O) for 7 weeks to saturate tissues with deuterium *in vivo*. Deuterium is rapidly incorporated throughout the body of an organism after treatment, bringing the deuterium enrichment in body water up to 5%. Deuterium enrichment is maintained through the intake of 8% ^2^H_2_O given as drinking water, thus making it an optimal labeling approach for a long-term *in vivo* experimental study. Mice were then given non-labeling drinking water to wash deuterium label out of faster generating tissues (i.e. the liver), but not enough time to significantly reduce label in slower lipid turn-over tissues such as the adipose. After 2 weeks, mice were injected intraperitoneally (IP) with either tunicamycin dissolved in DMSO, or DMSO control. Mice were treated with 1.5 mg/kg tunicamycin dissolved in DMSO, or DMSO control, and tissues were harvested 72 hours after the initial injection (n=4 mice per group).

### 4.4 Body water enrichment analysis

Mouse liver were distilled overnight upside down on a bead bath at 85°C to evaporate out body water. Deuterium present in the body water were exchanged onto acetone, and deuterium enrichment in the body water was measured via gas chromatography mass spectrometry (GC-MS)^36^.

### 4.5 Tissue preparation for liquid chromatography-mass spectrometry (LC-MS)

Tissues were flash frozen after harvest and homogenized in homogenization buffer (100 mM PMSF, 500 mM EDTA, EDTA-free Protease Inhibitor Cocktail (Roche, catalog number 11836170001), PBS) using a 5 mm stainless steel bead at 30 hertz for 45 seconds in a TissueLyser II (Qiagen). Samples were then centrifuged at 10,000 rcf for 10 minutes at 4°C. The supernatant was saved and protein was quantified using a Pierce BCA protein assay kit (ThermoFisher, catalog number 23225). 100 ug of protein was used per sample. 25 uL of 100 mM ammonium bicarbonate solution, 25 uL TFE, and 2.3 uL of 200 mM DTT were added to each sample and incubated at 60°C for 1 hour. 10 uL 200 mM iodoacetamide was then added to each sample and allowed to incubate at room temperature in the dark for 1 hour. 2 uL of 200 mM DTT was added and samples were incubated for 20 minutes in the dark. Each sample was then diluted with 300 uL H_2_O and 100 uL 100 mM ammonium bicarbonate solution. Trypsin was added at a ratio of 1:50 trypsin to protein (trypsin from porcine pancreas, Sigma Aldrich, catalog number T6567). Samples were incubated at 37°C overnight. The next day, 2 uL of formic acid was added. Samples were centrifuged at 10,000 rcf for 10 minutes, collecting the supernatant. Supernatant was speedvac’d until dry and re-suspended in 50 uL of 0.1 % formic acid/3% acetonitrile/96.9% LC-MS grade water and transferred to LC-MS vials to be analyzed via LC-MS.

### 4.6 Liquid chromatography-mass spectrometry (LC-MS) analysis

Trypsin-digested peptides were analyzed on a 6550 quadropole time of flight (Q-ToF) mass spectrometer equipped with Chip Cube nano ESI source (Agilent Technologies). High performance liquid chromatography (HPLC) separated the peptides using capillary and nano binary flow. Mobile phases were 95% acetonitrile/0.1% formic acid in LC-MS grade water. Peptides were eluted at 350 nl/minute flow rate with an 18 minute LC gradient. Each sample was analyzed once for protein/peptide identification in data-dependent MS/MS mode and once for peptide isotope analysis in MS mode. Acquired MS/MS spectra were extracted and searched using Spectrum Mill Proteomics Workbench software (Agilent Technologies) and a mouse protein database (www.uniprot.org). Search results were validated with a global false discovery rate of 1%. A filtered list of peptides was collapsed into a nonredundant peptide formula database containing peptide elemental composition, mass, and retention time. This was used to extract mass isotope abundances (M0-M3) of each peptide from MS-only acquisition files with Mass Hunter Qualitative Analysis software (Agilent Technologies). Mass isotopomer distribution analysis (MIDA) was used to calculate peptide elemental composition and curve-fit parameters for predicting peptide isotope enrichment based on precursor body water enrichment (p) and the number (n) of amino acid C-H positions per peptide actively incorporating hydrogen (H) and deuterium (D) from body water. Subsequent data handling was performed using python-based scripts, with input of precursor body water enrichment for each subject, to yield fractional synthesis rate (FSR) data at the protein level. FSR data were filtered to exclude protein measurements with fewer than 2 peptide isotope measurements per protein. Details of FSR calculations and data filtering criteria have been described in detail previously (Holmes et al., 2015).

### 4.7 Calculation of fractional replacement (f) and replacement rate constant (k) for individual proteins

Details of f calculations were previously described (Holmes et al., 2015).

### 4.8 Statistical analysis

Data were analyzed using GraphPad Prism software (version 8.0).

### 4.9 KEGG pathway analysis

Protein fractional synthesis rates were weighted by the peptide count and averaged according to their KEGG pathway involvements. We used the Uniprot.ws package in R from Bioconductor to find mappings between UniProt accession numbers and their corresponding KEGG IDs for each protein. Tables were generated for the entire known proteome for mouse. We then used the Bio.KEGG module of Biopython in Python to access to the REST API of the KEGG database to get a list of pathways to which each protein belongs. A set of all the pathways relevant to the experiment was generated and each protein and its corresponding fold change value were assigned to each pathway. KEGG pathways with no less than five proteins were used for representation of the data.

### 4.10 Tissue preparation for gas chromatography-mass spectrometry (GC-MS)

A chloroform methanol extraction was used to isolate lipids from the liver tissue. These lipids were run on a thin-layer chromatography (TLC) plate to separate phospholipid and triglyceride fractions. These fractions containing the palmitate were further derivatized for GC-MS analysis

### 4.11 Gas chromatography-mass spectrometry (GC-MS) analysis

Palmitate and cholesterol isotopic enrichments were measured by GC-MS (models 6890 and 5973; Hewlett-Packard, Palo Alto, CA) using a DB-225 fused silica column in methane chemical ionization mode monitoring m/z ratios of 159, 160, and 161 (representing M0, M1, and M2 isotopomers), as previously described (Hellerstein & Neese, 1997; Hellerstein, et al., 1991). Palmitate methyl ester enrichments were determined by GC-MS using a DB-17 column with helium as carrier gas and electron ionization mode monitoring m/z 270, 271, and 272 for M0, M1, and M2, as previously described^16^. Baseline, unenriched, standards were measured concurrently to correct for abundance sensitivity.

### 4.12 Calculation of *de novo* lipogenesis (DNL) and cholesterol synthesis

The measurement of newly synthesized fatty acids and total cholesterol formed during ^2^H_2_O labeling period was assessed using a combinatorial model of polymerization biosynthesis, as described previously (Hellerstein & Neese, 1997; Hellerstein, et al., 1991). Mass isotopomer distribution analysis (MIDA) is used to determine the number of hydrogen atoms in C–H bonds of fatty acids that were derived from cellular water during *de novo* synthesis of fatty acids, using body ^2^H_2_O to represent the precursor pool enrichment (p). Fractional and absolute contributions from DNL are then calculated. The value for f DNL represents the fraction of total triglyceride or phospholipid palmitate in the depot derived from DNL during the labeling period, and absolute DNL represents grams of palmitate synthesized by the DNL pathway.

### 4.13 RNAseq

RNA was isolated using standard Trizol protocol and RNA concentrations were obtained using a Nanodrop. Library preparation was performed using Kapa Biosystems mRNA Hyper Prep Kit. Sequencing was performed using NovaSeq, mode SR100 through the Vincent J. Coates Genomic Sequencing Core at University of California, Berkeley. Trimmed fastq reads were then aligned to the mouse genome and analyzed using Qiagen CLC Workbench Software. Differentially expressed genes were initially separated based on their direction (up/down). We then looked at which processes were enriched given the differentially expressed gene set with GOrilla^37^. We did a negative-log transform of the p-values for each significant enrichment and generated a figure using the Matplotlib package in Python.

### 4.14 Electron Microscopy

Electron microscopy was performed at the UC Berkeley Electron Microscope Laboratory. Samples were no larger than 0.5 mm and were agitated at each step. Samples were fixed for 1 hour in 2% glutaraldehyde in 0.1M sodium cacodylate buffer, pH 7.2, rinsed for 10 minutes in 0.1M sodium cacodylate buffer, pH 7.2 three times, put in 1% osmium tetroxide in 1.6% potassium ferricyanide for 1 hour, rinsed for 10 minutes in 0.1M sodium cacodylate buffer, pH 7.2, three times. Dehydrated in acetone: 35% acetone 10 minutes, 50% acetone 10 minutes, 70% acetone 10 minutes, 80% acetone 10 minutes, 95% acetone 10 minutes, 100% acetone 10 minutes, 100% acetone 10 minutes, 100% Acetone 10 minutes. Samples were then infiltrated with 2:1 acetone:resin (accelerator) for 1 hour, 1:1 acetone:resin for 1 hour, 75% acetone 25% resin overnight. The next morning, samples were put in pure resin for 1 hour, changed three times, and then pure resin plus accelerator for 1 hour. Samples were embedded into molds at 60◻ for 2 days with pure resin and accelerator. Samples were then visualized via the TECNAI 12 TEM.

### 4.15 Hematoxylin and eosin staining

Hematoxylin and eosin staining was performed at the UCSF Biorepository and Tissue. Images were collected with a Zeiss Plan-Apochromat 20x/0.8NA (WD=0.55mm) M27 Biomarker Technology Core. Imaging was conducted in a Zeiss Axio Scan.Z1 whole slide scanner objective lens in the brightfield mode with Hitachi HV-F202 camera.

## Acknowledgments

The authors would like to thank Marcy Matthews, Mark Fitch, and the entire Hellerstein lab for their technical assistance and feedback. We would also like to thank the Dillin lab at UC Berkeley for insightful input. Thanks also to Reena Zalpuri at the UC Berkeley Electron Microscope Laboratory for advice and assistance in electron microscopy sample preparation and data collection and to Monica Forsythe and all of the support staff at the UC Berkeley Animal Facility for assistance with mouse injections and animal care and housing. Graphical abstract and figures generated with BioRender. We thank Fred Ward for miscellaneous support.

## Author contributions

CPS, AD, and MKH conceived and designed experiments. CPS, LP, MC, SY, and RK performed experiments. LKJ performed RNA-seq assay and assisted with data analysis. HP assisted with tunicamycin treatment in mice. HM and EN assisted with LC-MS and GC-MS sample processing and data analysis. NZ wrote the script for KEGG pathway analysis. CPS, AEF, and MKH contributed to writing the manuscript.

## Supplementary Figures

**Supplement 1:**
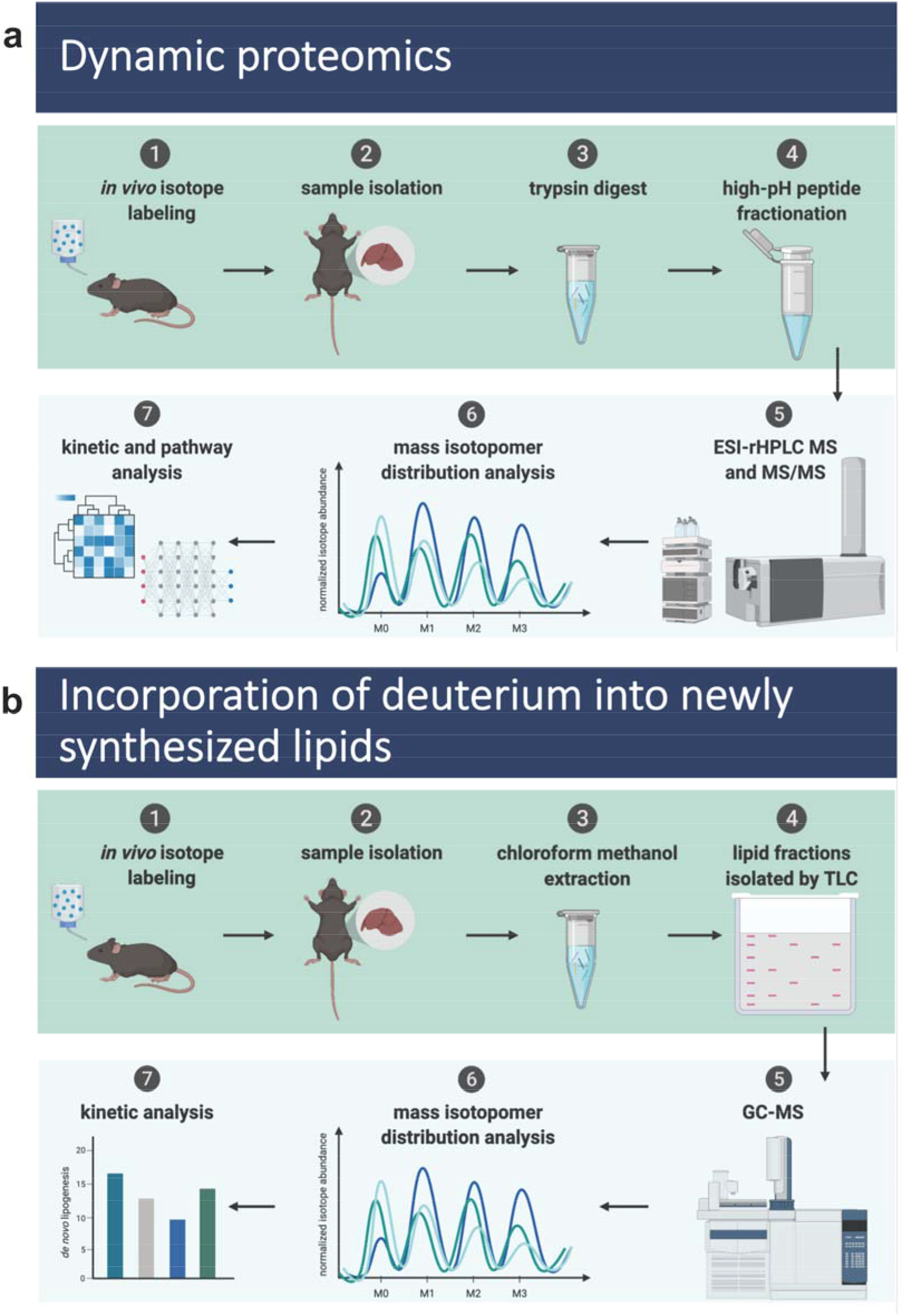
a) Experimental overview of dynamic proteomics approach. b) Experimental overview of measurement of *de novo* lipogenesis.

**Supplement 2:**
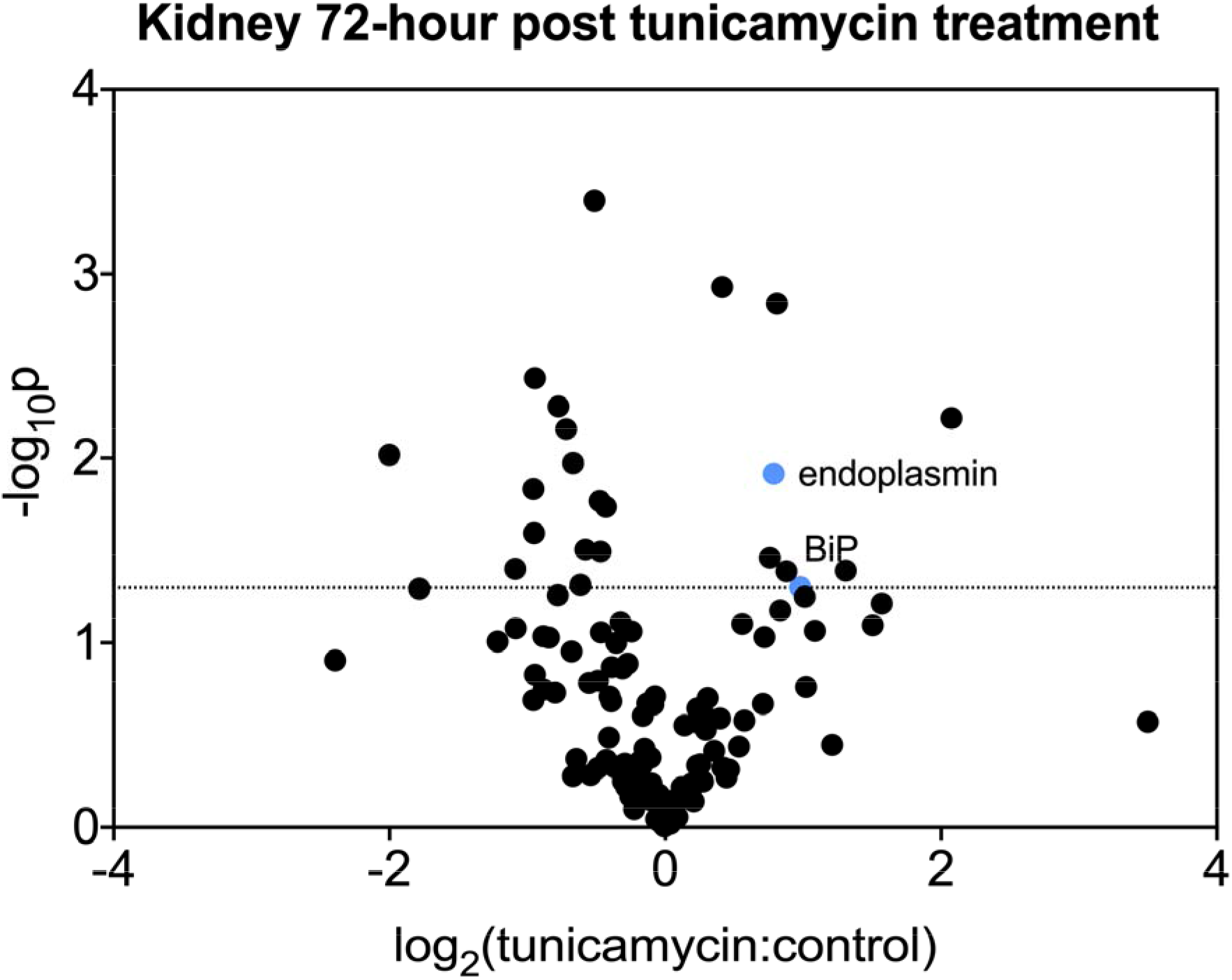
Ratio of tunicamycin:control treated mice kidneys. Volcano plot of all proteins for which fractional synthesis rates were measured 72 hrs post tunicamycin treatment. Points expressed as log2 fold-change tunicamycin treated/control on x-axis and − log10(p-value), obtained from 2-tail t-test, on y-axis.

## Notes

### Competing Interest Statement

The authors have declared no competing interest.

## References

1. Frakes, A. E. & Dillin, A. The UPRER: Sensor and Coordinator of Organismal Homeostasis. Mol. Cell 66, 761–771 (2017).

2. Wang, M. & Kaufman, R. J. Protein misfolding in the endoplasmic reticulum as a conduit to human disease. Nature 529, 326–335 (2016).

3. Salvadó, L., Palomer, X., Barroso, E. & Vázquez-Carrera, M. Targeting endoplasmic reticulum stress in insulin resistance. Trends Endocrinol. Metab. 26, 438–448 (2015).

4. Fu, S., Watkins, S. M. & Hotamisligil, G. S. The role of endoplasmic reticulum in hepatic lipid homeostasis and stress signaling. Cell Metab. 15, 623–634 (2012).

5. Cnop, M., Foufelle, F. & Velloso, L. A. Endoplasmic reticulum stress, obesity and diabetes. Trends Mol. Med. 18, 59–68 (2012).

6. Rutkowski, D. T. et al. UPR pathways combine to prevent hepatic steatosis caused by ER stress-mediated suppression of transcriptional master regulators. Dev. Cell 15, 829–840 (2008).

7. Rutkowski, D. T. Liver function and dysfunction - a unique window into the physiological reach of ER stress and the unfolded protein response. FEBS J. 286, 356–378 (2019).

8. Hotamisligil, G. S. Endoplasmic reticulum stress and the inflammatory basis of metabolic disease. Cell 140, 900–917 (2010).

9. Bertolotti, A., Zhang, Y., Hendershot, L. M., Harding, H. P. & Ron, D. Dynamic interaction of BiP and ER stress transducers in the unfolded-protein response. Nat. Cell Biol. 2, 326–332 (2000).

10. Walter, P. & Ron, D. The Unfolded Protein Response: From Stress Pathway to Homeostatic Regulation. Science vol. 334 1081–1086 (2011).

11. Lee, J.-S., Mendez, R., Heng, H. H., Yang, Z.-Q. & Zhang, K. Pharmacological ER stress promotes hepatic lipogenesis and lipid droplet formation. Am. J. Transl. Res. 4, 102–113 (2012).

12. DeZwaan-McCabe, D. et al. ER Stress Inhibits Liver Fatty Acid Oxidation while Unmitigated Stress Leads to Anorexia-Induced Lipolysis and Both Liver and Kidney Steatosis. Cell Rep. 19, 1794–1806 (2017).

13. Herrema, H. et al. XBP1s Is an Anti-lipogenic Protein. J. Biol. Chem. 291, 17394–17404 (2016).

14. Schuck, S., Prinz, W. A., Thorn, K. S., Voss, C. & Walter, P. Membrane expansion alleviates endoplasmic reticulum stress independently of the unfolded protein response. J. Cell Biol. 187, 525–536 (2009).

15. Holmes, W. E., Angel, T. E., Li, K. W. & Hellerstein, M. K. Dynamic Proteomics: In Vivo Proteome-Wide Measurement of Protein Kinetics Using Metabolic Labeling. Methods Enzymol. 561, 219–276 (2015).

16. Neese, R. A. et al. Measurement of endogenous synthesis of plasma cholesterol in rats and humans using MIDA. Am. J. Physiol. 264, E136–47 (1993).

17. Turner, S. M. et al. Measurement of TG synthesis and turnover in vivo by 2H2O incorporation into the glycerol moiety and application of MIDA. Am. J. Physiol. Endocrinol. Metab. 285, E790–803 (2003).

18. Mohan, S., R, P. R. M., Brown, L., Ayyappan, P. & G, R. K. Endoplasmic reticulum stress: A master regulator of metabolic syndrome. Eur. J. Pharmacol. 860, 172553 (2019).

19. Ron, D. & Walter, P. Signal integration in the endoplasmic reticulum unfolded protein response. Nat. Rev. Mol. Cell Biol. 8, 519–529 (2007).

20. Rendleman, J. et al. New insights into the cellular temporal response to proteostatic stress. Elife 7, e39054 (2018).

21. Reid, D. W., Chen, Q., Tay, A. S. L., Shenolikar, S. & Nicchitta, C. V. The unfolded protein response triggers selective mRNA release from the endoplasmic reticulum. Cell (2014).

22. Shankaran, M. et al. Circulating protein synthesis rates reveal skeletal muscle proteome dynamics. J. Clin. Invest. 126, 288–302 (2016).

23. Greenbaum, D., Colangelo, C., Williams, K. & Gerstein, M. Comparing protein abundance and mRNA expression levels on a genomic scale. Genome Biol. 4, 117 (2003).

24. Liu, Y., Beyer, A. & Aebersold, R. On the dependency of cellular protein levels on mRNA abundance. Cell (2016).

25. Maier, T., Güell, M. & Serrano, L. Correlation of mRNA and protein in complex biological samples. FEBS Lett. 583, 3966–3973 (2009).

26. Vandewynckel, Y.-P. et al. Modulation of the unfolded protein response impedes tumor cell adaptation to proteotoxic stress: a PERK for hepatocellular carcinoma therapy. Hepatol. Int. 9, 93–104 (2015).

27. Borgese, N., Francolini, M. & Snapp, E. Endoplasmic reticulum architecture: structures in flux. Curr. Opin. Cell Biol. 18, 358–364 (2006).

28. Daniele, J. R. et al. UPRER promotes lipophagy independent of chaperones to extend life span. Sci Adv 6, eaaz1441 (2020).

29. Sriburi, R., Jackowski, S., Mori, K. & Brewer, J. W. XBP1: a link between the unfolded protein response, lipid biosynthesis, and biogenesis of the endoplasmic reticulum. J. Cell Biol. 167, 35–41 (2004).

30. Colgan, S. M., Tang, D., Werstuck, G. H. & Austin, R. C. Endoplasmic reticulum stress causes the activation of sterol regulatory element binding protein-2. Int. J. Biochem. Cell Biol. 39, 1843–1851 (2007).

31. Han, J. & Kaufman, R. J. The role of ER stress in lipid metabolism and lipotoxicity. J. Lipid Res. 57, 1329–1338 (2016).

32. Ho, N. et al. Stress sensor Ire1 deploys a divergent transcriptional program in response to lipid bilayer stress. J. Cell Biol. 219, (2020).

33. Estrada, A. L. et al. Short-term changes in diet composition do not affect in vivo hepatic protein synthesis in rats. Am. J. Physiol. Endocrinol. Metab. 314, E241–E250 (2018).

34. Fu, S. et al. Polysome profiling in liver identifies dynamic regulation of endoplasmic reticulum translatome by obesity and fasting. PLoS Genet. 8, e1002902 (2012).

35. Smith, G. I., Mittendorfer, B. & Klein, S. Metabolically healthy obesity: facts and fantasies. J. Clin. Invest. 129, 3978–3989 (2019).

36. Yang, D. et al. Assay of Low Deuterium Enrichment of Water by Isotopic Exchange with [U-13C3]Acetone and Gas Chromatography–Mass Spectrometry. Analytical Biochemistry vol. 258 315–321 (1998).

37. Eden, E., Navon, R., Steinfeld, I., Lipson, D. & Yakhini, Z. GOrilla: a tool for discovery and visualization of enriched GO terms in ranked gene lists. BMC Bioinformatics 10, 48 (2009).

